# Hyperactivity of the Amygdala Mediates Depressive-Like Phenotypes and Decreased Serotonin Release

**DOI:** 10.1101/2025.03.10.642393

**Authors:** Kyuhee Kim, Yuanying Lai, Qianqian Gao, Huajie Wang, Kwok Tsz Ying, Muhammad Asim, Jufang He

## Abstract

Clinical and preclinical studies have consistently demonstrated a correlation between hyperactivity of the amygdala and the onset of depression. However, the underlying mechanisms influencing serotonin levels a critical neurotransmitter implicated in depression and a primary target for selective serotonin reuptake inhibitors (SSRIs) remain inadequately understood. In this study, we employed a restrained inescapable shock (RIS) model to investigate these mechanisms in mice. The RIS paradigm elicited depressive-like phenotypes, increased c-Fos expression in the amygdala, diminished serotonin levels, and elevated corticosterone concentrations. Notably, chemogenetic inhibition of the amygdala mitigated depressive symptoms, reduced neuronal activity in this region, and restored serotonin levels. Anatomical analyses revealed a significant connectivity between the central amygdala (CeA) and the dorsal raphe nucleus (DRN). Fiber photometry recordings indicated that serotonergic neuronal activity in the DRN decreased in response to aversive stimuli, accompanied by amygdala activation following RIS, with no notable alterations in DRN GABAergic activity. These findings suggest that chronic stress may exacerbate amygdala hyperactivity, which subsequently inhibits serotonin release in the brain, potentially intensifying depressive states. Therefore, targeting amygdala hyperactivity may represent a novel therapeutic strategy for the management of stress-related depressive and anxiety disorders.

## Introduction

Major depressive disorder (MDD) is a severe mental health condition that affects approximately 280 million people worldwide. According to a scientific report from the World Health Organization (WHO), the global prevalence of anxiety and depression increased by 25% during the first year of the COVID-19 pandemic (Santomauro et al., 2021). As prescriptions for selective serotonin reuptake inhibitors (SSRIs) such as fluoxetine, sertraline, and paroxetine have risen dramatically, many patients continue to experience low efficacy and various side effects (Bartova et al., 2019; Sanacora et al., 2012). This highlights an urgent need to understand how stress-induced alterations in neural circuits contribute to the development of mood disorders.

Stress is recognized as a major risk factor for the onset of mood disorders, including depressive disorders (Asim, Wang, Chen, et al., 2024; Liu et al., 2020). The amygdala often exhibits hyperactivity in individuals with mood disorders, and this stress-induced hyperactivity in the amygdala may represent a critical mechanistic pathway linking chronic stress exposure to the emergence of depressive symptoms and related psychopathology (Nikolova et al., 2018). Recent studies in rodents have shown that chronic stress increases glutamatergic neuronal activity and decreases specific subtypes of GABAergic neuronal activity in the amygdala while enhancing inhibition through high-frequency activation of GABAergic neurons in the basolateral amygdala (BLA) alleviates stress-mediated depressive-like behaviors (Asim, Wang, Chen, et al., 2024; Asim, Wang, Waris, et al., 2024). However, the impact of amygdala hyperactivity on the serotonin system, a primary target of current antidepressant drugs such as fluoxetine and sertraline, remains unclear.

The dorsal raphe nucleus (DRN) is a primary source of serotonin (5-HT) and projects to various brain regions (Lesch & Waider, 2012; Ren et al., 2019). Previous studies have reported a decrease in serotonin release in chronically stressed rodent models (Nazzi et al., 2019; Prakash et al., 2020) and lower serotonin levels in depressed human subjects (Kostanjšak & Zdunić, 2017; Obermanns et al., 2021). Moreover, antidepressants such as fluoxetine and sertraline are known to maintain elevated synaptic serotonin levels by inhibiting the serotonin transporter (Pawluski et al., 2020; Talaee et al., 2024). Therefore, understanding the dynamic interactions between the amygdala and the DRN a key serotonergic hub has important implications for elucidating the neural mechanisms underlying stress-related mood disorders, altered serotonin release, and developing more effective treatments.

In this study, we first establish a RIS mouse model that induces depressive-like behaviors along with altered cortisol and serotonin levels. We then employ a chemogenetic approach to inhibit the amygdala to determine if this intervention can alleviate depressive-like behaviors and attenuate alterations in cortisol and serotonin levels in the stressed model. Next, we use retrograde and anterograde tracing techniques to investigate whether the DRN, a serotonin-enriched region, receives inputs from the amygdala. Finally, we adopt fiber photometry to explore changes in general and cell-type-specific neuronal activity in the DRN in response to acute footshock aversive stimuli paired with amygdala activation before and after the RIS procedure. Through this series of experiments, we highlight that RIS leads to decreased serotonin neuronal activity and release, implying that amygdala hyperactivity during stress and depression alters serotonin release. While inhibition of the amygdala could represent a potential therapeutic strategy to address stress-related depressive-like behaviors.

## Material & Methods

### Animals

All experimental protocols were reviewed and authorized by the Animal Subject Ethics Sub-Committee of the City University of Hong Kong. In this study, we utilized male C57BL/6N mice aged 8 to 10 weeks for behavioral assessments, immunohistochemistry, fiber photometry, optogenetics, and anatomical investigations. These mice, along with Ai14 (B6;129S6-Gt(ROSA)26Sortm14(CAG-tdTomato)Hze/J, C57 background) mice, were sourced from The Jackson Laboratory. The mice were housed in social groups of five within cages that were maintained in a temperature-regulated room at 23±1°C. A 12-hour light/dark cycle was followed, with lights on from 8:00 AM to 8:00 PM. The mice had unrestricted access to food and water. To ensure consistency, all behavioral tests were conducted during the animals’ active nighttime period. All possible measures were taken to reduce the number of animals used and to alleviate their suffering.

### Restrained inescapable shock stress paradigm

Each group of C57BL/6 animals was exposed to a 3-day stress protocol and compared to a control group that was subjected to gentle handling. Mice were 8 weeks old when the RIS stress protocol was initiated. Each group consisted of 10 male C57BL/6 mice. Four days prior to the start of the experiments, the mice were separated and housed individually. They were randomly assigned to either experimental or control groups and kept in ventilated cages with bedding, food, and water. The mice remained in the same cages for the duration of the experiment. On Day 1, the mice were placed in a restraint chamber and subjected to one hour of restraint stress, during which they received 30-120 electric shocks (20 to 30 seconds apart for 3 to 5 seconds each, with intensities ranging from 0.2 mA to 1 mA, and mice received 30 electric shocks from 0.7 mA to 1mA). The shocks began at 0.2 mA and were gradually increased by 0.1 mA, with the protocols conducted at 2 PM. On Day 2, the same procedures were followed, and after completion, the mice were kept in deprivation until the next day. On Day 3, the mice returned to the restraint chamber and received 30 electric shocks of varying intensities (0.2, 0.4, 0.8, and 1.0 mA), each lasting a maximum of 30 seconds. These procedures took place at 8 AM. All the electric shocks were delivered within 60 minutes.

### Behavior test

Mice were transported to the testing room and allowed to habituate for 30 minutes before the behavioral tests. Behavioral assessments were conducted in the following sequence: Sucrose preference test (SPT), Open field test (OFT), and Tail suspension test (TST). All behavioral tests were conducted under dim illumination. The instruments were thoroughly cleaned with 70% ethanol between animals to eliminate olfactory cues.

### Sucrose preference test (SPT)

Mice were habituated to a two-bottle choice for 48 hours containing tap water and then one bottle was switched to a 1% sucrose solution on testing day. They had access to both bottles, and the bottle positions were switched every 2 hours to avoid place preference. The sucrose preference ratio was determined by dividing the total consumption of the sucrose solution by the total consumption of both the water and sucrose solutions combined.

### Open field test (OFT)

The open field test assesses the balance between exploring a new environment and a rodent’s natural fear of open spaces. Mice individually explored an open field arena (50 × 50 × 40 cm) for 10 minutes. Their locomotor activity and time spent in the central(30 × 30 cm) and peripheral zones were automatically tracked and quantified using Smart 3.0 software.

### Tail suspension test (TST)

The tail-suspension test assesses various interventions expected to influence depression-related behaviors. In this procedure, mice are suspended by their tails using tape, preventing them from escaping or grasping onto nearby surfaces. Mobility was measured over a 6-minute period using a Smart 3.0 software system.

### Hormone Measurements

Blood was collected via tail venipuncture after each mouse was removed from its home cage and kept at room temperature for 2 hours. The serum was then separated by centrifugation at 13,000 g for 5 minutes at 4°C and stored at −80°C. Serum cortisol and serotonin levels were measured using enzyme-linked immunosorbent assay (ELISA) kits from Abcam.

### Viruses

The viruses utilized are as follows: AAV9-CamKiia-hM4Di-mCherry-WPRE-PA (5.33 × 1012 vg/ml), AAV9-TPH2-Cre (5.29 × 1012 vg/ml), AAV9-mDlx-GCaMP6s-WPRE-SV40 (1.00 × 1013 vg/ml) was obtained from BrainVTA company (Wuhan, China). pAAV-hSyn-EGFP (3.30×1013 vg/ml), pENN-AAV/retro-hSyn-Cre-WPRE-hGH (1.90×1013 vg/ml), pAAV9-hSyn-DIO-mCherry (2.30×1013 vg/ml), pENN-AAV1-hSyn-Cre-WPRE-hGH (1.90×1013 vg/ml), pAAV-Syn-ChrimsonR-tdT (2.60×1013 vg/ml), pAAV-Syn-GCaMP6s-WPRE-SV40 (2.70×1013 vg/ml), pAAV-hsyn-hM4D(Gi)-mCherry (2.00×1013 vg/ml), pAAV-Syn-Flex-GCaMP6s-WPRE-SV40 (2.50×1013 vg/ml) were purchased from Addgene (MA, USA).

### Stereotactic surgery

For the stereotactic surgeries, the mice were first anesthetized by administering sodium pentobarbital (50mg/kg) via intraperitoneal injection. Viruses were bilaterally injected in BLA, CeA, or DRN. A small opening was made in the skull using a dental drill to precisely target the desired brain regions. The stereotactic coordinates of injection sites were determined regarding the mouse brain atlas. Following this, the appropriate viral vectors were slowly infused into the target brain regions using a glass pipette attached to the Hamilton syringe at a rate of 50nl/min to minimize tissue damage. The injection coordinates were as follows: BLA (anteroposterior [AP] =−1.60mm, mediolateral [ML]=±3.37mm, dorsoventral [DV] =−4.00mm from bregma and dura mater). CeA (anteroposterior [AP] =−1.25mm, mediolateral [ML]=±3.00mm, dorsoventral [DV] =−4.6mm from bregma). DRN (anteroposterior [AP] =−4.3mm, mediolateral [ML]=1.00mm, dorsoventral [DV] =−2.6mm at 16° from lateral to medial) A total volume of 200-300nl of the virus was infused into each brain region. After the infusion, the injection needle was left in place for an additional 5 minutes before it was slowly withdrawn to allow for complete diffusion of the virus. For fiber photometry, optical fibers (4mm length) were carefully implanted above DRN (AP: −4.3mm, ML: 1.00mm, DV: −2.5mm at 16° from lateral to medial). For optogenetic stimulation, mice were implanted with bilateral optical fiber cannulas (6 mm length) held in ceramic ferrules (Inper company) over the CeA (anteroposterior [AP] =−1.25mm, mediolateral [ML]=±3.00mm, dorsoventral [DV] =−4.45mm from bregma and dura matter). The surgical site was then closed using dental cement, and the animals were given sufficient time, at least 3 weeks, to recover from the surgery before any further experimentation was conducted.

### Fiber photometry

We performed fiber photometry to figure out how DRN neural activity will be changed during footshock (aversive stimuli) and footshock paired with CeA activation. Mice were allowed to habituate to the fiber photometry enclosure for 30 minutes before the start of the experiment. For footshock, mice received 3 trials of foot shocks (0.2 mA for 2s) separated by a 1-minute inter-trial interval. For footshock paired with laser stimulation of CeA, mice received 3 trials of footshock, each immediately preceded by 4 seconds of laser stimulation (40Hz) over the CeA. The fluorescent signals emitted by the GCaMP6s indicator within the DRN were continuously monitored throughout these experiments using a fiber photometry system, providing real-time insights into neural activity within this brain region in response to the different stimuli. The fitted 405-nm signal was used to normalize the 473-nm signal with the following formula: ΔF/F0 = (473-nm signal - fitted 405-nm signal) / fitted 405-nm signal. This method of normalization allows for the determination of relative changes in fluorescence (ΔF/F0) by comparing the 473-nm signal to the fitted 405-nm signal.

### Immunohistochemistry

Mice were deeply anesthetized, then intracardially perfused with phosphate-buffered saline (PBS) followed by 4% paraformaldehyde (PFA) in PBS, 60min after the stress procedures. The brains were carefully removed and fixed in 4% PFA overnight. Coronal sections, each 50 μm thick, were prepared using a vibratome. The sections were washed with PBS and blocked for 2 hours with 5% goat serum in PBST (0.3% Triton X-100 in PBS) to minimize non-specific binding. They were then incubated overnight at 4°C with primary antibodies, including anti-c-fos rabbit antibody (ab190289; 1:1000), anti-5-HT1A antibody (ab85), and anti-GAD67 antibody (ab2938; 1:200), to enable specific detection of proteins. After washing the sections three times with PBS, they were incubated with secondary antibodies, Alexa Fluor 488 (1:1000) and Alexa Fluor 594 (1:500), for 2 hours at room temperature to visualize the target proteins. Following another three washes with PBS, the slices were stained with DAPI (1:10000) for 5 minutes to label the cell nuclei and subsequently washed three times with PBS. The sections were then mounted onto slides, and images were captured using a Nikon confocal microscope to visualize the labeled proteins and nuclei. Data analysis was performed using NIS Elements software to analyze the images and extract relevant information. Cell counts were conducted throughout the BLA, CeA, and DRN, and the averages were calculated from slices of 3-5 mice.

### Chemogenetic manipulation

To investigate the effect of chemogenetic inhibition on the BLA in alleviating depression-like behavior in our stress model, we bilaterally injected pAAV-CamKii-hM4Di-mCherry or pAAV-CamKii-mCherry into the BLA of mice. Three weeks post-injection, the mice received CNO (5 mg/kg) for three consecutive days to activate the inhibitory hM4Di receptor before undergoing behavioral tests. We injected pAAV-hsyn-hM4Di-tdTomato or AAV-hsyn-mCherry into the CeA to investigate how chemogenetic silencing of the CeA affects stress-induced depression-like phenotypes in the CeA-DRN circuit. Subsequently, mice were implanted with optical fibers or cannulas in the DRN to facilitate targeted manipulations of neural activity in these regions.

### Quantification and Statistical Analyses

Data are presented as means ± s.e.m. Statistical analyses were conducted using GraphPad Prism 8.0.1 software and Origin Pro 2024. Paired and unpaired t-tests were conducted to compare two groups with normally distributed data. For multiple groups, one-way and repeated measure two-way analysis of variance (ANOVA) was employed, followed by the Bonferroni test for post hoc comparisons. Behavioral data were analyzed without knowledge of group assignments. Statistical significance was indicated as *P < 0.05,* *P < 0.01, ***P < 0.001, and ****P < 0.0001.

## Result

### RIS-induced depressive-like behavior accompanied by changes in BLA c-fos activity and level of corticosterone and serotonin

In this study, we aimed to investigate the impact of the hyperactivity of the amygdala on serotonin activity levels and depressive-like phenotypes. To accomplish this, we first utilized a RIS paradigm to induce depressive-like phenotypes in mice. After RIS exposure, we conducted behavioral tests, including the open field test, tail suspension test, and sucrose preference test, to assess the consequences of RIS. Moreover, we also performed a follow-up behavioral experiment 6 weeks later to evaluate the long-term behavioral effects of the RIS (**Figure 1A**). In the sucrose preference test, RIS mice showed significantly less preference for sucrose compared to control mice, indicating the development of anhedonia, a core symptom of major depression (**Figure 1B**). Even 6 weeks after the RIS model, sucrose preference remained diminished. The immobility observed in the tail suspension test was significantly greater in RIS mice than in controls, reflecting a depression-like phenotype (**Figure 1C**). Furthermore, the immobility time remained higher in RIS mice compared to controls even after 6 weeks. In the open field test, RIS mice displayed reduced exploratory behavior, indicated by less time spent in the center of the arena, suggesting increased anxiety-like behavior (**Figure 1D-F**).

**Figure 1.**
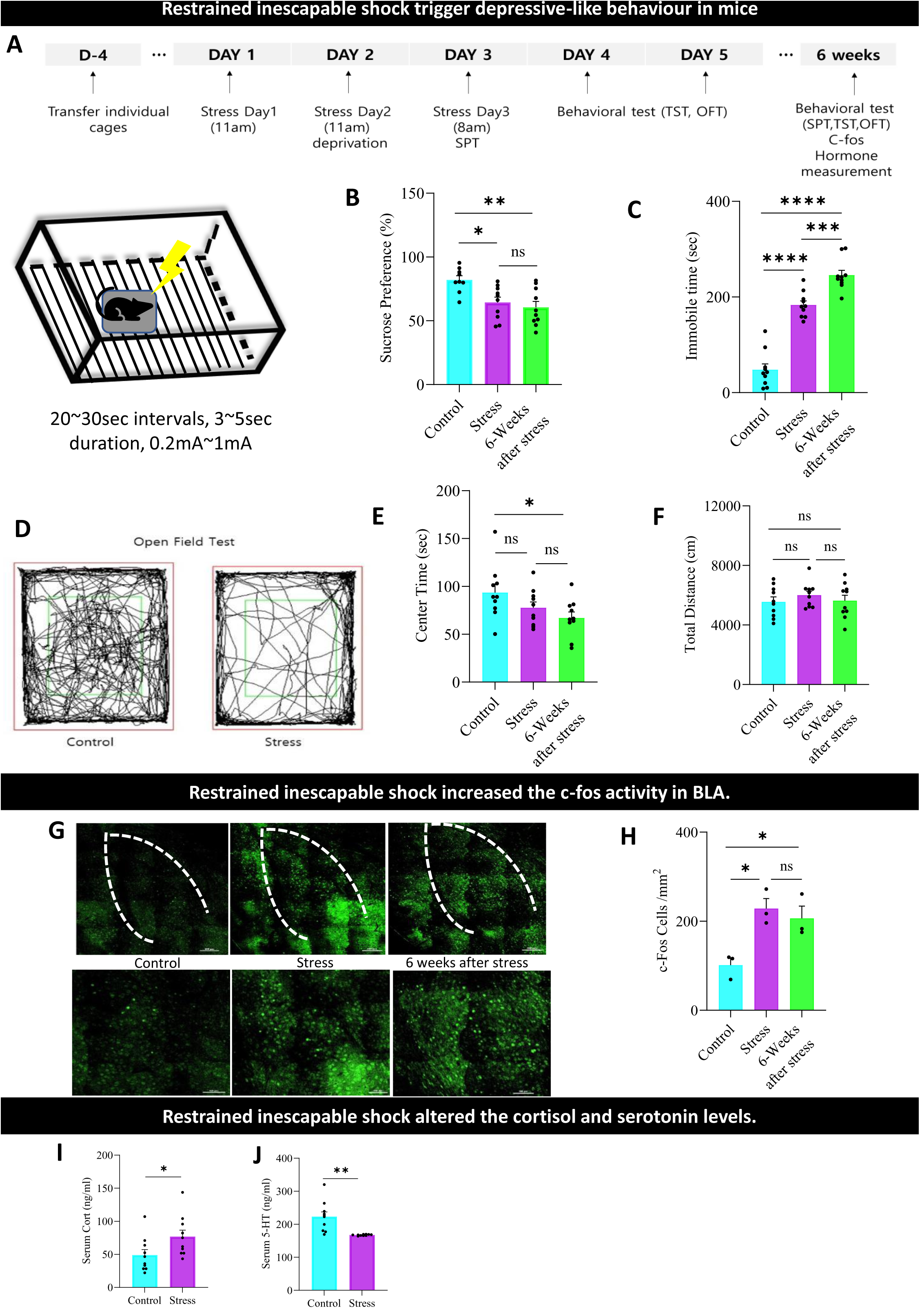
Restrained inescapable shock triggers depressive-like behavior in mice. **A.** An Experimental design to induce depressive-like behavior in rodents. **B.** Sucrose preference percentage between control, stress, and 6 weeks after stress group. One-way ANOVA, Treatment (column): **P < 0.0023, F (2, 26) = 7.721. Control vs. Stress: *P< 0.0157, t = 3.049, df = 26. Control vs. 6weeks after stress: **P < 0.0030, t = 3.712, df = 26. Stress Vs. 6weeks after stress: ^ns^P > 0.9999, t = 0.6817, df = 26. Adjustment: Bonferroni. N = 9-10 mice. **C**. Immobility time (sec) during TST. One-way ANOVA, Treatment (column): ****P < 0.0001, F (2, 27) = 96.53. Control vs. Stress: ****P< 0.0001, t = 9.310, df = 27. Control vs. 6weeks after stress: ****P < 0.0001, t = 13.59, df = 27. Stress Vs. 6weeks after stress: ***P <0.0006, t = 4.277, df = 27. Adjustment: Bonferroni. N = 10 mice. **D.** Trajectory map during OFT. **E.** Time spent in center (sec) during OFT. One-way ANOVA, Treatment (column): *P < 0.0464, F (2, 27) = 3.448. Control vs. Stress: ^ns^P< 0.3883, t = 1.564, df = 27. Control vs. 6weeks after stress: *P < 0.0439, t = 2.609, df = 27. Stress Vs. 6weeks after stress: ^ns^P <0.9163, t = 1.045, df = 27. Adjustment: Bonferroni. N = 10 mice. **F.** Total distance explored (cm) during OFT. One-way ANOVA, Treatment (column): ^ns^P < 0.5752, F (2, 27) = 0.5646. Control vs. Stress: ^ns^P< 0.9883, t = 0.9932, df = 27. Control vs. 6weeks after stress: ^ns^P > 0.9999, t = 0.1695, df = 27. Stress Vs. 6weeks after stress: ^ns^P > 0.9999, t = 0.8237, df = 27. Adjustment: Bonferroni. N = 10 mice. **G.** c-fos expression control (Left), stress (Middle), and 6 weeks after stress (Right) group in BLA. **H.** Measured c fos expression levels in BLA. One-way ANOVA, Treatment (column): *P < 0.0155, F (2, 6) = 9.044. Control vs. Stress: *P< 0.0219, t = 3.979, df = 6. Control vs. 6weeks after stress: *P < 0.0499, t = 3.290, df = 6. Stress Vs. 6weeks after stress: ^ns^P > 0.9999, t = 0.6893, df = 6. Adjustment: Bonferroni. N = 3 mice. **I.** Showing the corticosterone level changes between control and depressive-like behavior mice. Unpaired t-test, *P < 0.0421, t = 2.188, df = 18, N = 10 mice. **J.** Showing the 5-ht level changes between control and depressive-like behavior mice. Unpaired t-test, **P < 0.0012, t = 3.856, df = 18, N = 10 mice.

Our previous study showed that social defeat stress increased c-fos activity in the BLA (M. Asim et al., 2023). Therefore, we examined if RIS could also increase c-fos activity in the BLA. Indeed, we found that the RIS mice exhibited significantly elevated c-fos activity in the BLA compared to controls (**Figure 1G-H**), suggesting heightened neural activity in this region. Previous studies suggest that chronic stress is known to lead to dysregulation of the hypothalamic-pituitary-adrenal axis leading to altered corticosterone and serotonin (5-HT) levels along with depression-like behaviors (Hutton et al., 2015; Rosenkranz et al., 2010). To further explore the physiological underpinnings of the observed depressive-like phenotype in our RIS model, we measured the levels of glucocorticoids and 5-HT, two key neuromodulators implicated in the pathogenesis of depression. Our results revealed a significant elevation in the plasma concentration of corticosterone, the primary glucocorticoid in rodents, in RIS mice compared to their non-stressed counterparts (**Figure 1I**). Importantly, our analysis also demonstrated that RIS mice exhibited markedly reduced levels of serotonin (**Figure 1J**), a critical neurotransmitter for mood regulation, within the amygdala compared to control animals. Our findings demonstrate that the chronic stress paradigm used in our study reliably induces a persistent depressive-like phenotype in mice along with altered corticosterone and 5-HT level.

### Chemogenetic inhibition of the BLA reduced depressive-like behavior, linked to decreased c-fos activity, lower corticosterone, and higher serotonin levels

In light of the observed increase in BLA c-fos activity in the RIS model, we sought to investigate whether selectively inhibiting BLA neurons could alleviate depressive-like phenotypes in the RIS model. We employed a chemogenetic approach to selectively silence the BLA in the RIS model by injecting pAAV-CaMKIIa-hM4Di-mCherry or the control virus pAAV-CaMKIIa-mCherry into the BLA. This allowed us to reversibly inhibit BLA neurons upon administration of the synthetic ligand clozapine-N-oxide (CNO). Three weeks after viral vector injections, mice underwent the RIS paradigm. Behavioral testing was conducted with pharmacological inhibition of the BLA achieved by intraperitoneal injection of CNO 30 minutes before each test (**Figure 2 A-B**).

**Figure 2.**
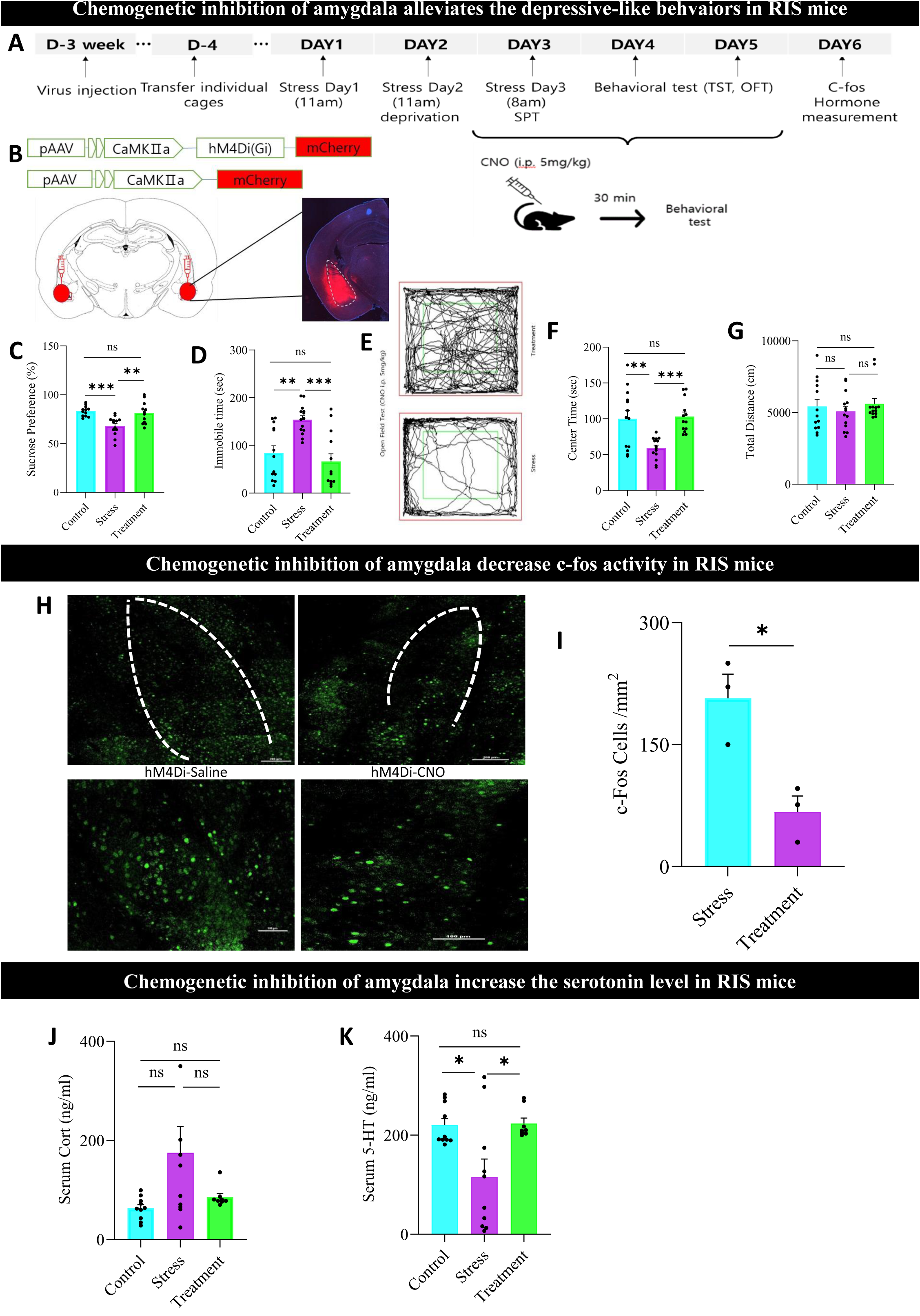
Chemogenetic inhibition of the amygdala alleviates depressive-like behaviors in RIS mice. **A.** Schematical representation of experimental design. **B.** hM4Di-mCherry virus expression for chemogenetic inhibition of BLA. **C.** Sucrose preference percentage among control, stress, and hM4Di group. One-way ANOVA, Treatment (column): ***P < 0.0003, F (2, 34) = 10.52. Control vs. Stress: ***P< 0.0006, t = 4.189, df = 34. Control vs. Treatment: ^ns^P > 0.9999, t = 0.5268, df = 34. Stress Vs. Treatment: **P < 0.0026, t = 3.652, df = 34. Adjustment: Bonferroni. N = 13-14 mice. **D.** Immobility time (sec) during TST. One-way ANOVA, Treatment (column): ****P < 0.0001, F (2, 37) = 12.04. Control vs. Stress: **P< 0.021, t = 3.692, df = 37. Control vs. Treatment: ^ns^P > 0.9999, t = 0.9030, df = 37. Stress Vs. Treatment: ***P <0.0001, t = 4.612, df = 37. Adjustment: Bonferroni. N = 13-14 mice. **E.** Trajectory map during OFT. **F.** Time spent in center (sec) during OFT. One-way ANOVA, Treatment (column): ***P < 0.0003, F (2, 37) = 10.12. Control vs. Stress: **P< 0.0020, t = 3.721, df = 37. Control vs. Treatment: ^ns^P > 0.9999, t = 0.2835, df = 37. Stress Vs. Treatment ***P < 0.0009, t = 4.010, df = 37. Adjustment: Bonferroni. N = 13-14 mice. **G.** Total distance explored(cm) during OFT. One-way ANOVA, Treatment (column): ^ns^P < 0.4281, F (2, 37) = 0.6486. Control vs. Stress: ^ns^P > 0.9999, t = 0.5915, df = 37. Control vs. Treatment: ^ns^P > 0.9999, t = 0.3233, df = 37. Stress Vs. Treatment ^ns^P < 0.9999, t = 0.9207, df = 37. Adjustment: Bonferroni. N = 13-14 mice. **H.** c-fos expression among control, stress, and hM4Di group in BLA. **I.** Measured c-fos expression levels in BLA. Unpaired t-test, *P < 0.0171, t = 3.928, df = 4, N = 3 mice. **J.** Showing the corticosterone level changes among control, stress, and hM4Di groups. One-way ANOVA, Treatment (column): ^ns^P < 0.0530, F (2, 25) = 3.311. Control vs. Stress: ^ns^P < 0.0646, t = 2.452, df = 25. Control vs. Treatment: ^ns^P > 0.9999, t = 0.4684, df = 25. Stress Vs. Treatment: ^ns^P > 0.2313, t = 1.844, df = 25. Adjustment: Bonferroni. N = 8-10 mice. **K.** The 5-ht level changes control, stress, and the hM4Di group. One-way ANOVA, Treatment (column): **P < 0.0057, F (2, 25) = 6.410. Control vs. Stress: *P < 0.0137, t = 3.116, df = 25. Control vs. Treatment: ^ns^P > 0.9999, t = 0.08747, df = 25. Stress Vs. Treatment: *P < 0.0171, t = 3.025, df = 25. Adjustment: Bonferroni, N = 8-10 mice.

The sucrose preference test revealed that chemogenetic inhibition in stressed mice expressing hM4Di showed a significant increase in sucrose consumption, rescuing the anhedonic phenotype (**Figure 2 C**). Importantly, CNO administration also reduced immobility time in the tail suspension test for stressed mice expressing hM4Di, indicating an antidepressant-like effect (**Figure 2 D**). In the open field test, chemogenetic inhibition of the BLA in stressed mice restored their time spent in the center of the arena to levels comparable to non-stressed controls, indicating reduced anxiety-like behavior (**Figure 2 E-G**). Additionally, CNO-treated stressed mice expressing hM4Di exhibited decreased c-Fos activity in the BLA compared to stressed control mice, confirming the efficacy of the chemogenetic inhibition **(Figure 2 H-I).** To further understand the neurochemical mechanisms underlying the observed behavioral effects of BLA inhibition, we measured the corticosterone and serotonin levels. We found that the RIS-induced decrease in serotonin levels was effectively reversed by chemogenetic silencing of the BLA **(Figure 2 K).** However, no significant alterations in cortisol levels were observed in the treatment group **(Figure 2 J)**. Our findings suggest that targeted inhibition of the BLA could represent a promising therapeutic strategy to alleviate depressive-like behaviors.

### Retrograde and anterograde tracing confirmed the connectivity between the amygdala and the dorsal raphe

Given the increased activity in the amygdala and decreased serotonin levels, we aimed to investigate whether the amygdala projects to the DRN, a region known for supplying serotonin to various brain areas, potentially leading to reduced serotonin levels. We confirmed the anatomical connectivity between the amygdala and the DRN. To achieve this, we first injected the retrograde pENN-AAV/Retro-hSyn-Cre-WPRE-hGH virus into DRN Ai14 mice (**Figure 3A**). This experimental setup enabled visualization of retrograde tdTomato expression in structures sending input to the DRN. Interestingly, we did not observe retrograde tdTomato-positive cells in the BLA; however, abundant tdTomato expression was seen in the central amygdala (CeA) (**Figure 3 B).** We further validated the CeA-DRN connectivity using two distinct anterograde labeling methods (**Figure 3 A & D**). In the first set of mice, we injected AAV-hSyn-EGFP into the CeA, confirming fiber expression in the DRN (**Figure 3 C**). In the second set, we injected AAV1-hSyn-Cre into the CeA and AAV-hSyn-DIO-mCherry into the DRN, allowing us to express the mCherry reporter in DRN neurons receiving input from the CeA (**Figure 3 D-E)**. These retrograde and anterograde labeling approaches collectively confirmed the anatomical connectivity between the CeA and DRN. It is well established that glutamatergic neurons in the BLA project to the CeA (Babaev et al., 2018; Hartley et al., 2019; Namburi et al., 2015), which is predominantly composed of GABAergic neurons (comprising approximately 95% of total cells) (Marek et al., 2013). This suggests that hyperactivity in the BLA may enhance the activity of GABAergic neurons in the CeA, which in turn project to serotonergic neurons in the DRN, thereby inhibiting serotonin release in the brain, as observed during depressive states (Cleare, 1997; Obermanns et al., 2021). Moreover, we confirmed the subtypes of DRN neurons receiving input from the amygdala by utilizing GABA and 5-HT antibodies in slices where mCherry was expressed in DRN cells receiving input from the CeA. We found that 55.5 % of the mCherry cells were colocalized with 5-HT cells, while 45.5% of the mCherry cells were also colocalized with GAD-67 cells (**Figure 3 F-H**), confirming the input from the CeA to DRN serotonin-producing neurons. These anatomical findings suggest that hyperactivity of the amygdala during depression may inhibit serotonin release through GABAergic inputs from the CeA to the DRN.

**Figure 3.**
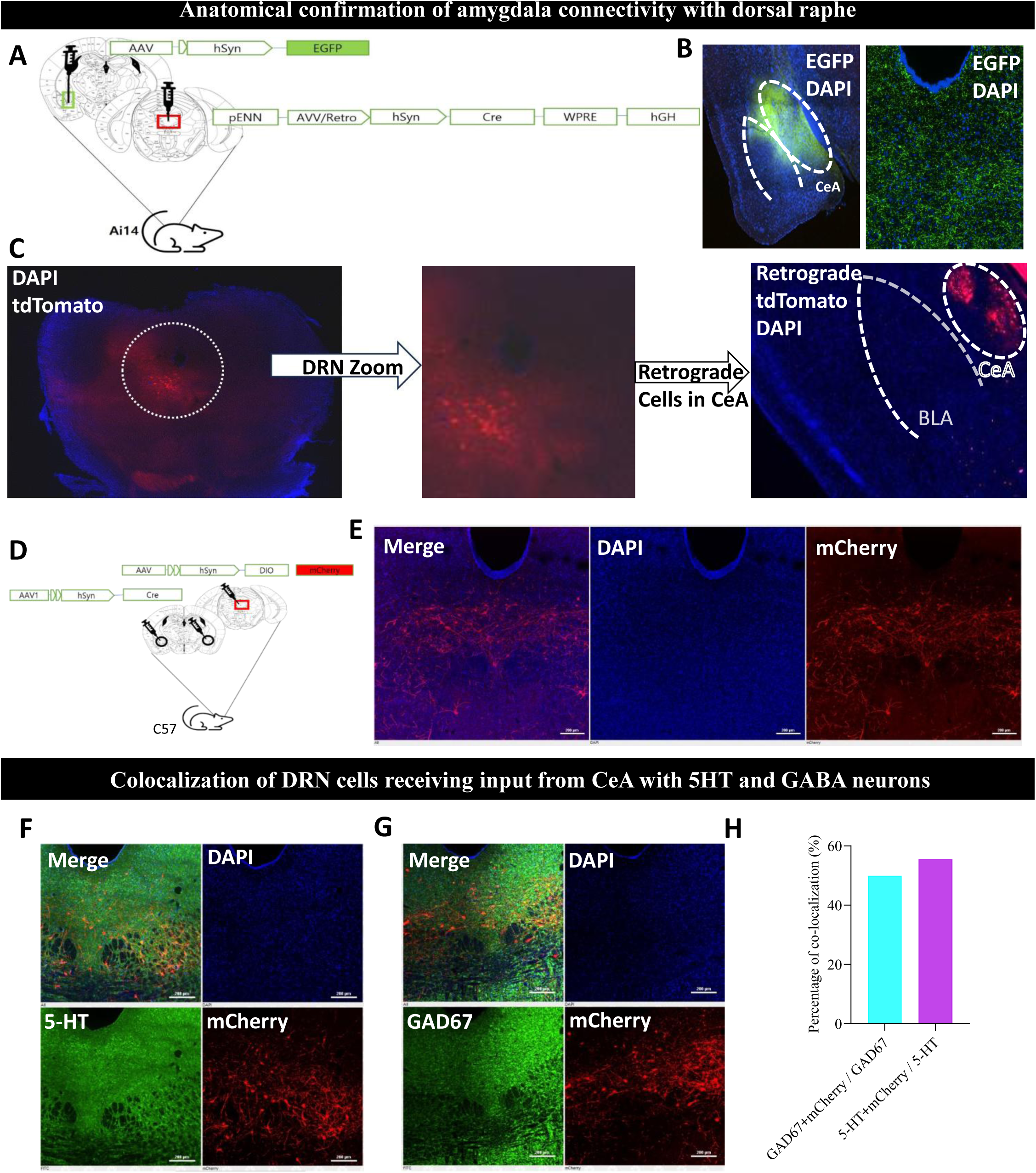
Anatomical confirmation of the amygdala connectivity with dorsal raphe nucleus using a retrograde virus. **A.** Schematical representation of the experimental design for AAV-retro virus injection in DRN. **B.** EGFP expression in CeA and fibers in DRN. **C.** tdTtomato expression in DRN Ai 14 mice, and tdTomato regrade cells in CeA. **D.** Schematical representation of the experimental design for Cre-dependant mCherry expression in CeA-DRN pathway. **E.** mCherry expression in DRN. **F.** Co-localization with 5-HT antibody. **G.** Co-localization with GAD67 positive-cells. **H**. Percentage of co-localized cells with 5HT and GAD-67 in DRN.

### DRN neuronal activity in response to aversive stimuli and CeA activation before and after the RIS depression model

Next, we investigated whether the RIS depression model induced changes in the neuronal activity of the DRN. To do this, we recorded fiber photometry calcium responses in the DRN of freely moving mice during footshock stimuli, both alone and paired with optogenetic activation of the CeA, before and after RIS. Mice were injected with AAV-Syn-ChrimsonR-tdTomato into the CeA and pAAV-Syn-GCamp6S-WPRE-SV40 into the DRN (**Figure 4 A-C**). Before the RIS model, exposure to footshocks resulted in a significant increase in calcium transients in the DRN, indicating increased neural activity in response to the acute aversive stimulus (**Figure 4 D-E**). This finding underscores the sensitivity of DRN neurons to acute stress and suggests their vital role in processing aversive experiences. However, we did not observe any significant changes in neural activity following the RIS model (**Figure 4 D-G**). Additionally, there were no differences in DRN neuronal activity in response to footshock versus footshock paired with CeA activation before the RIS model (**Supplementary Figure S1 A-C**). Interestingly, when mice were subjected to footshock paired with optogenetic activation of the CeA, DRN neuronal activity exhibited a slight increase after the RIS model compared to before (**Figure 4 H-K**). In contrast to our hypothesis, activation of the CeA enhances overall DRN neuronal activity following the RIS model when paired with footshock. This observation underscores a more complex interaction between these two brain regions.

**Figure 4.**
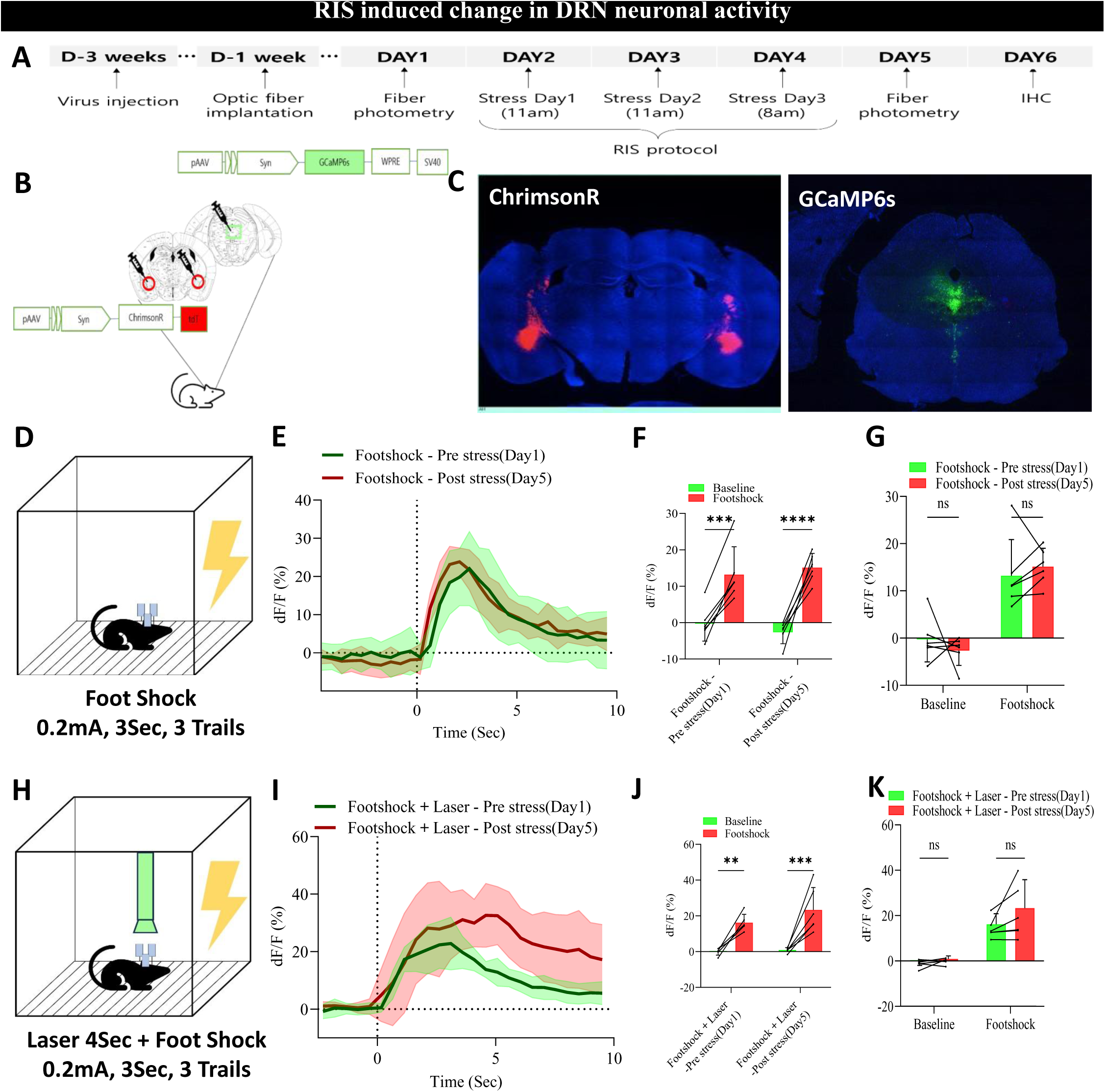
DRN neuronal activities in response to Foot shock and Foot shock paired with CeA activation. **A.** An Experimental Design. **B.** Schematical representation of experimental design for Fiber photometry. **C.** ChrimsonR virus expression in CeA and GCaMP6s virus expression in the DRN. **D.** Schematical representation of the experimental design for foot shock. **E.** Showing calcium traces during FS in experimental mice with restrained inescapable shock. **F.** Average change in DRN activity before and during FS in experimental mice with restrained inescapable shock. RM Two-way ANOVA, Interaction: ^ns^P < 0.19, F (1, 10) = 2.018. Row Factor: ^ns^P < 0.93, F (1, 10) = 0.008. Column Factor: ****P < 0.0001, F (1, 10) = 109.7. Baseline vs FS (Pre-Stress-Day1): ***P < 0.0002, t = 6.402, df = 10. Baseline vs FS (Post Stress-Day5): ****P < 0.0001, t = 8.411, df = 10. Adjustment: Bonferroni. N = 6 mice. **G.** Showing the average changes during FS in experimental mice with restrained inescapable shock. RM Two-way ANOVA, Interaction: ^ns^P < 0.31, F (1, 10) = 1.104. Row Factor ****P < 0.0001, F (1, 10) = 51.92. Column Factor: ^ns^P < 0.91, F (1, 10) = 0.013. Baseline (Pre-Stress-Day1) vs Baseline (Post Stress-Day5): ^ns^P > 0.86, t = 0.82, df = 10. FS (Pre-Stress-Day1) vs FS (Post Stress-Day5): ^ns^P < 0.99, t = 0.66, df = 10. Adjustment: Bonferroni. N = 6 mice. **H.** Schematical representation of the experimental design for foot shock paired with CeA activation. **I.** Showing calcium traces during FS paired with CeA activation in experimental mice with restrained inescapable shock. **J.** Average change in DRN activity before and during FS paired with CeA activation in experimental mice with restrained inescapable shock. RM Two-way ANOVA, Interaction: ^ns^P < 0.27, F (1, 10) = 1.36. Row Factor: nsP < 0.19, F (1, 10) = 1.93. Column Factor: ****P < 0.0001, F (1, 10) = 52.72. Baseline vs FS-Laser (Pre-Stress-Day1): **P < 0.0031, t = 4.31, df = 10. Baseline vs FS-Laser (Post Stress-Day5): ***P < 0.0003, t = 5.96, df = 10. Adjustment: Bonferroni. N = 6 mice. **K.** Showing the average changes during FS paired with CeA activation in experimental mice with restrained inescapable shock. RM Two-way ANOVA, Interaction: ^ns^P < 0.15, F (1, 10) = 2.33. Row Factor: ***P < 0.0002, F (1, 10) = 33.05. Column Factor: nsP < 0.076, F (1, 10) = 3.90. Baseline (Pre-Stress-Day1) vs Baseline (Post Stress-Day5): ^ns^P < 0.99, t = 0.32, df = 10. FS-Laser (Pre-Stress-Day1) vs FS-Laser (Post Stress-Day5): nsP < 0.066, t = 2.48, df = 10. Adjustment: Bonferroni, N = 6 mice.

Given the complex interaction between the CeA and DRN, we aimed to investigate the activity of specific neuronal populations within the DRN that may be differentially altered before and after the RIS model. First, we injected pAAV-Syn-ChrimsonR-tdTomato into the CeA and pAAV-Syn-mDlx-GCamp6s-WPRE-SV40 into the DRN, selectively labeling GABAergic neurons with GCaMP6s (**Figure 5 A-C**). Our results demonstrated that the activity of GABAergic neurons significantly increased in response to footshock; however, there was no significant difference in neuronal activity observed before and after the RIS model (**Figure 5 D-F**). Furthermore, we did not detect any differences in neuronal activity between footshock and footshock paired with CeA activation prior to the RIS model (**Supplementary Figure S2 A-D**). Additionally, there was no change in GABAergic neuronal activity in response to footshock paired with CeA activation before and after the RIS model (**Figure 5 G-I**). Although we did not observe any differences in neuronal activity pre- and post-RIS depression model, the activation of GABAergic neurons in response to footshock stimuli suggests that these neurons may also play a role in modulating local circuits and serotonin release.

**Figure 5.**
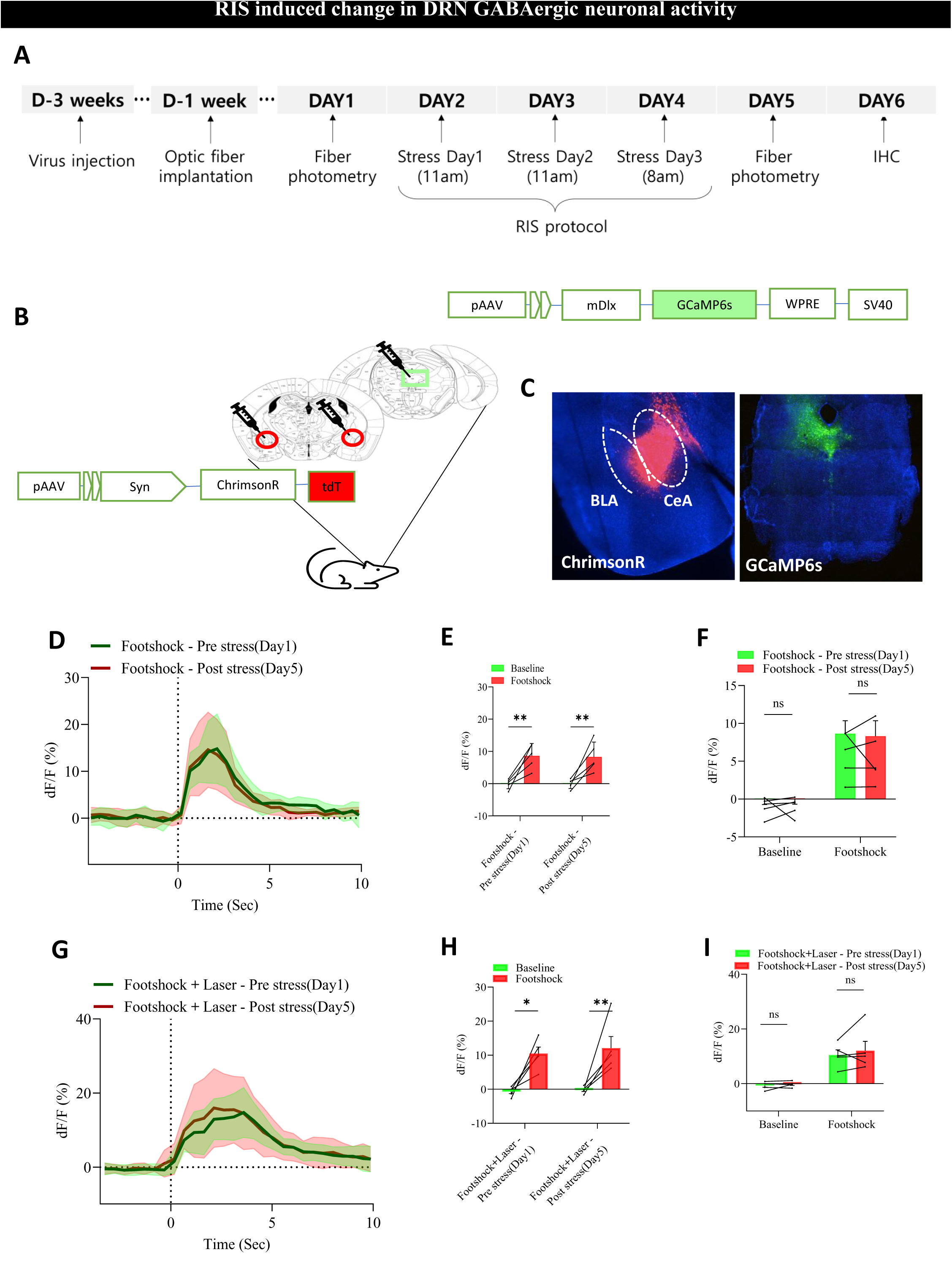
RIS induced a change in DRN GABAergic neuronal activity. **A.** An experimental design. **B.** Schematical representation of the experimental design for GCaMP6s virus injection. **C.** ChrimsonR virus expression in CeA and GCaMP6s virus expression in the DRN. **D.** Showing calcium traces during FS in mDlx-GCaMP6s virus expression mice with restrained inescapable shock. **E.** Average change in DRN activity before and during FS in mDlx-GCaMP6s virus expression mice with restrained inescapable shock. RM Two-way ANOVA, Interaction: ^ns^P < 0.91, F (1, 8) = 0.015. Row Factor: ^ns^P < 0.91, F (1, 8) = 0.012. Column Factor: ***P < 0.0004, F (1, 8) = 34.40. Baseline vs FS (Pre-Stress-Day1): **P < 0.0057, t = 4.23, df = 8. Baseline vs FS (Post Stress-Day5): **P < 0.0073, t = 4.061, df = 8. Adjustment: Bonferroni. N = 5 mice. **F.** Showing the average changes between FS in mDlx-GCaMP6s virus expression mice with restrained inescapable shock. RM Two-way ANOVA, Interaction: ^ns^P < 0.85, F (1, 8) = 0.039. Row Factor: ****P < 0.0015 F (1, 8) = 22.12. Column Factor: ^ns^P < 0.87, F (1, 8) = 0.030. Baseline (Pre-Stress-Day1) vs Baseline (Post Stress-Day5): ^ns^P > 0.9999, t = 0.016, df = 8. FS (Pre-Stress-Day1) vs FS (Post Stress-Day5): ^ns^P > 0.9999, t = 0.26, df = 8. Adjustment: Bonferroni. N = 5 mice. **G.** Showing calcium traces during FS paired with CeA activation in mDlx-GCaMP6svirus expression mice with restrained inescapable shock. **H.** Average change in DRN activity before and during FS paired with CeA activation in mDlx-GCaMP6s virus expression mice with restrained inescapable shock. RM Two-way ANOVA, Interaction: ^ns^P < 0.79, F (1, 8) = 0.07. Row Factor: ^ns^P < 0.59, F (1, 8) = 0.30. Column Factor: ***P < 0.0005, F (1, 8) = 31.30. Baseline vs FS-Laser (Pre-Stress Day1): *P < 0.011, t = 3.77, df =8. Baseline vs FS-Laser (Post Stress-Day5): **P < 0.0065, t = 4.15, df = 8. Adjustment: Bonferroni. N = 5 mice. **I.** The average changes during FS paired with CeA activation in mDlx-GCaMP6s virus expression mice with restrained inescapable shock. RM Two-way ANOVA, Interaction: ^ns^P < 0.64, F (1, 8) = 0.63. Row Factor: ***P < 0.002, F (1, 8) = 2081. Column Factor: ^ns^P < 0.39, F (1, 8) = 0.81. Baseline (Pre-Stress-Day1) vs Baseline (Post Stress-Day5): ^ns^P > 0.9999, t = 0.29, df = 8. FS-Laser (Pre-Stress-Day1) vs FS Laser (Post Stress-Day5): ^ns^P > 0.71, t = 0.98, df = 8. Adjustment: Bonferroni, N = 5 mice.

We next injected AAV9-TPH2-Cre and pAAV-Syn-FLEX-GCamp6s-WPRE-SV40 into the DRN, along with AAV-Syn-ChrimsonR-tdTomato into the CeA, to selectively record serotonergic neurons in the DRN in response to footshock or CeA activation paired with footshock, both before and after the RIS model (**Figure 6 A-B**). Our findings indicated a slight elevation in the activity of DRN serotonergic neurons in response to acute footshock compared to baseline activity. However, we observed a decreased DRN serotonergic neuronal activity in response to footshock when comparing pre-and post-RIS model conditions (**Figure 6 D & F**). Interestingly, when we paired footshock with CeA activation, we noted a slight trend to increased serotonergic neuronal activity compared to footshock alone before the RIS model (**Supplementary Figure S2 E-H**). However, we found that footshock paired with activation of CeA neurons led to decreased activation of DRN serotonergic neuronal activity after RIS compared to before (**Figure 6 G-I**). These results suggest that while GABAergic neuronal activity did not significantly change in response to the RIS depression model during CeA activation, the activity of serotonergic neurons decreased. These insights pave the way for further exploration of the intricate dynamics between these regions, which could inform therapeutic strategies for addressing stress-related disorders.

**Figure 6.**
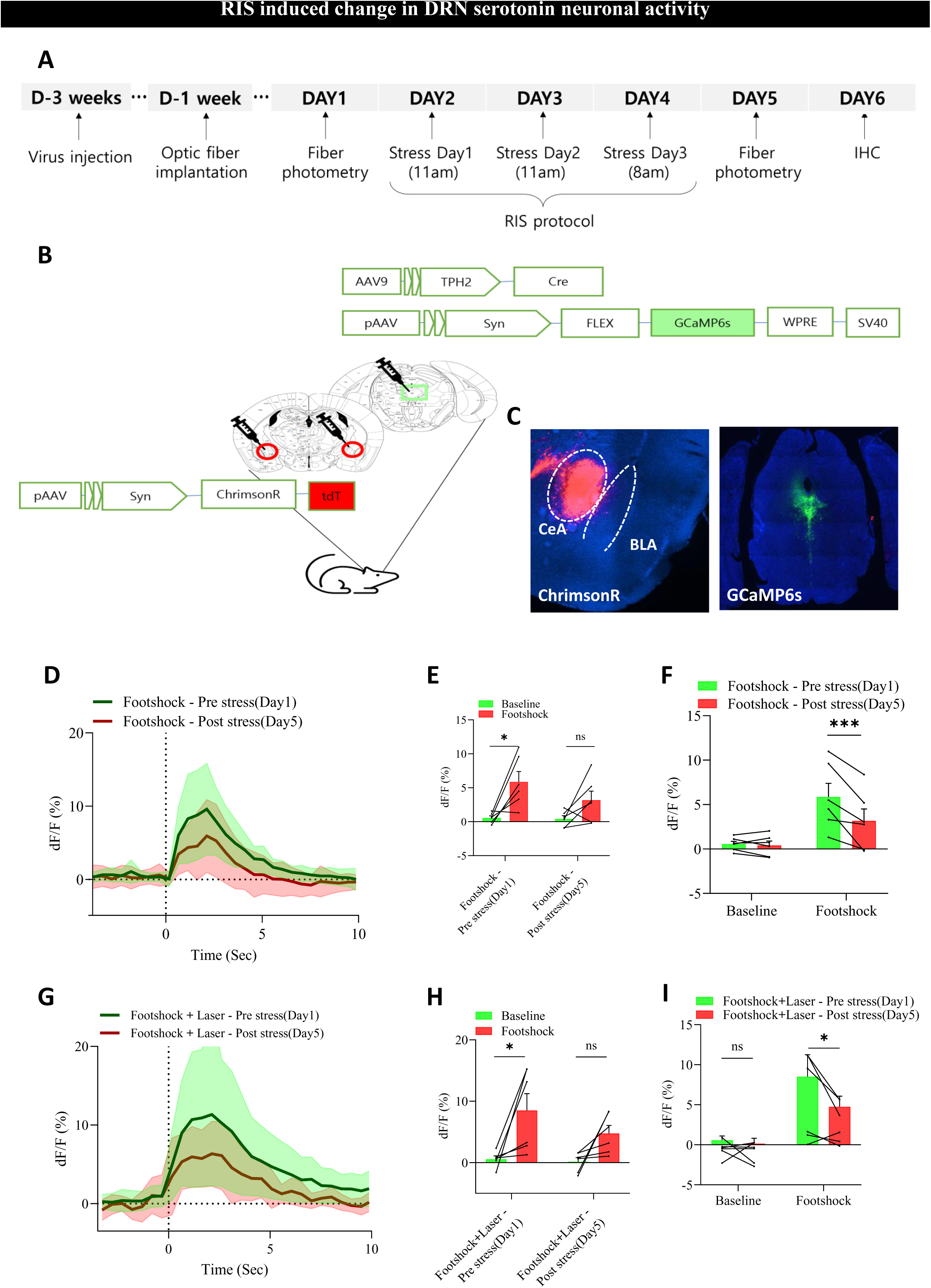
RIS induced a change in DRN serotonin neuronal activity. **A.** An experimental design. **B.** Schematical representation of the experimental design for GCaMP6s virus injection. **C.** ChrimsonR virus expression in CeA and GCAMP6s virus expression in the DRN. **D.** Showing calcium traces during FS in TPH2-GCaMP6s expression mice with restrained inescapable shock. **E.** Average change in DRN activity before and during FS in TPH2-GCaMP6s expression mice with restrained inescapable shock. RM Two-way ANOVA, Interaction: ^ns^P < 0.26, F (1, 10) = 1.44. Row Factor: ^ns^P < 0.20, F (1, 10) = 1.83. Column Factor: **P < 0.0033, F (1, 10) = 14.69. Baseline vs FS (Pre-Stress-Day1): *P < 0.0103, t = 3.56, df = 10. Baseline vs FS (Post Stress-Day5): ^ns^P < 0.18, t = 1.86, df = 10. Adjustment: Bonferroni. N = 6 mice. **F.** Showing the average changes between FS in TPH2 GCaMP6s expression mice with restrained inescapable shock. RM Two-way ANOVA Interaction: **P < 0.006, F (1, 10) = 12.44. Row Factor: *P < 0.02, F (1, 10) = 7.86 Column Factor: **P < 0.003, F (1, 10) = 15.49. Baseline (Pre-Stress Day1) vs Baseline (Post Stress-Day5): ^ns^P > 0.9999, t = 0.29, df = 10. FS (Pre-Stress-Day1) vs FS (Post Stress-Day5): ***P < 0.0007, t = 5.28, df = 10. Adjustment: Bonferroni. N = 6 mice. **G.** Showing calcium traces during FS paired with CeA activation in TPH2-GCaMP6s expression mice with restrained inescapable shock. **H.** Average change in DRN activity before and during FS paired with CeA activation in TPH2-GCaMP6s expression mice with restrained inescapable shock. RM Two-way ANOVA, Interaction: ^ns^P < 0.35, F (1, 10) = 0.94. Row Factor: nsP < 0.17, F (1, 10) = 2.13 Column Factor: **P < 0.0044, F (1, 10) = 13.38. Baseline vs FS-Laser (Pre-Stress-Day1): *P < 0.016, t = 3.28, df = 10. Baseline vs FS-Laser (Post Stress-Day5): nsP < 0.17, t = 1.89, df = 10. Adjustment: Bonferroni. N = 6 mice. **I.** The average changes during FS paired with CeA activation in TPH2-GCaMP6s expression mice with restrained inescapable shock were shown. RM Two-way ANOVA, Interaction: ^ns^P < 0.12, F (1, 10) = 0.122. Row Factor: *P < 0.0106, F (1, 10) = 9.82. Column Factor: nsP < 0.0609, F (1, 10) = 4.46. Baseline (Pre-Stress-Day1) vs Baseline (Post Stress-Day5): ^ns^P < 0.99, t = 0.29, df = 10. FS-Laser (Pre-Stress-Day1) vs FS Laser (Post Stress-Day5): *P < 0.046, t = 2.69, df = 10. Adjustment: Bonferroni. N = 6 mice.

## Discussions

Clinical (Groenewold et al., 2013; Qiao et al., 2020) and preclinical studies (Asim, Wang, Chen, et al., 2024; Asim, Wang, Waris, et al., 2024; Rosenkranz et al., 2010) have consistently reported that hyperactivity of the amygdala is associated with depression. However, the mechanisms by which amygdala hyperactivity affects serotonin levels an important neurotransmitter implicated in depression and a primary target for current antidepressants, such as SSRIs (Clevenger et al., 2018; Dale et al., 2016) remain unclear. In this study, we found that the RIS model induced depressive-like phenotypes in mice, characterized by increased amygdala activity and decreased serotonin levels, alongside elevated corticosterone levels, consistent with previous findings (Muhammad Asim et al., 2023; Moncrieff et al., 2023; Pretorius, 2004). Notably, chemogenetic inhibition of the amygdala not only alleviated depressive symptoms but also reduced neuronal activity in this region and improved serotonin levels. Anatomical investigations confirmed the connectivity between the CeA and the DRN, a region rich in serotonergic neurons (Paquelet et al., 2022). Furthermore, fiber photometry recordings indicated that serotonergic neuronal activity in the DRN decreased in response to aversive stimuli when coupled with amygdala activation after the RIS, with no significant alterations observed in GABAergic neuronal activity within the DRN. This suggests that persistent stress may lead to increased amygdala activity, which subsequently inhibits serotonin release in the brain. This inhibition could diminish the positive effects of serotonin (Heifets et al., 2019; Walsh et al., 2021) and exacerbate negative environmental stressors, resulting in depressive-like states. Therefore, targeting the hyperactivity of the amygdala may represent a promising strategy for managing stress-related depressive and anxiety-like states.

One major challenge in rodent models of depression is the prolonged duration of stress protocols required to induce depressive phenotypes, such as the chronic unpredictable stress model (4-8 weeks) (Bijata et al., 2022), chronic restraint model (2-3 weeks) (Son et al., 2019), and social defeat (10 days) (Golden et al., 2011), with many animals failing to exhibit depressive behaviors (Cerniauskas et al., 2019; Planchez et al., 2019). In this study, we employed a 3-day restraint stress combined with tail shock, successfully establishing a long-lasting model of depressive phenotypes observed even 6 weeks later. This short-term stress protocol is as effective as longer methods, offering significant time and labor savings. While the learned helplessness model also induces depressive-like phenotypes with 3 days of inescapable foot shock, it primarily focuses on escape latency, neglecting other critical phenotypes like anhedonia and despair (Abdulla et al., 2024; Chourbaji et al., 2005). Therefore, our 3-day restraint stress with tail shock protocol presents a more reliable and efficient approach to induce multiple depressive phenotypes.

Consistent with previous studies (Asim, Wang, Chen, et al., 2024; Asim, Wang, Waris, et al., 2024; Rosenkranz et al., 2010), we found that RIS induced amygdala hyperactivity, which may serve as a potential biomarker for depressive states, as several studies have reported amygdala hyperactivity in depressed patients (Boukezzi et al., 2022; Groenewold et al., 2013; Qiao et al., 2020). Additionally, prior research has indicated decreased serotonin and increased corticosterone levels in depressive states (Muhammad Asim et al., 2023; Moncrieff et al., 2023; Pretorius, 2004). Our findings further demonstrated that RIS elevated corticosterone and reduced serotonin, reinforcing the notion that these hormonal changes are associated with depression. Notably, chemogenetic inhibition of the amygdala not only alleviated depressive-like symptoms but also reversed the levels of serotonin. This suggests that targeting amygdala hyperactivity could have therapeutic implications for treating depression and supports the hypothesis that decreased amygdala activity correlates with lower stress levels (Runia et al., 2023; Sudimac et al., 2022). Moreover, the hyperactivity of the amygdala could be a potential source for decreased serotonin levels observed in depressed patients (Kostanjšak & Zdunić, 2017; Obermanns et al., 2021).

Anatomical investigations revealed that the CeA exhibits dense projections to the DRN. The CeA is composed of approximately 95% GABAergic neurons (Marek et al., 2013), which are activated by the BLA (Babaev et al., 2018; Hartley et al., 2019; Namburi et al., 2015), containing glutamatergic neurons as a major population. This suggests that GABAergic projections from the CeA may inhibit serotonin neurons in the DRN, potentially leading to decreased serotonin levels in the brain during depressive states. Indeed, our findings indicate that CeA inhibition of serotonin neuronal activity in the DRN occurs following RIS when paired with footshock, without any concomitant change in GABAergic activity within the DRN. Interestingly, we observed a slight increase in the overall activity of DRN neurons post-RIS, suggesting the presence of a subpopulation of DRN neurons that may be activated by the CeA, potentially through a disinhibition mechanism (Babaev et al., 2018; Ye & Veinante, 2019). This could occur if some interneurons within the CeA inhibit local GABAergic projection neurons. Future studies are warranted to elucidate the underlying mechanisms in greater detail, particularly to determine whether specific subsets of CeA neurons are inhibited during depressive states, if these neurons project to the DRN, and what types of cells (GABAergic or serotonergic) they target. A deeper understanding of the cell-type-specific connectivity between the CeA and DRN is essential for advancing our knowledge in this area.

In sum, this study demonstrates that the RIS model induces depressive-like states in mice, characterized by altered hormone levels and increased amygdala activity. The findings suggest that this effect may be mediated by increased GABAergic inhibition from the CeA to the serotonergic neurons in the DRN. Furthermore, the inhibition of the amygdala appears to alleviate both the depressive symptoms and the dysregulation of hormone levels, highlighting a potential target for therapeutic intervention in mood disorders.

## Acknowledgments

This work was supported by funding from the following: Hong Kong Research Grants Council, General Research Fund: CityUHK 11101521, CityUHK 11103922, CityUHK 11104923, and CityUHK 11104524. Hong Kong Research Grants Council, Collaborative Research Fund: C1043-21G. Hong Kong Research Grants Council, Theme-Based Research Scheme: T13-605/18-W. Hong Kong Research Grants Council, Senior Research Fellow Scheme: SRFS2324-1S02. Innovation and Technology Fund of the Hong Kong SAR, China: GHP_075_19GD. Hong Kong Health Bureau, Health and Medical Research Fund: 09203656, 08194106. Innovation Technology Commission of the Hong Kong SAR, China: Health@InnoHK program. We also thank the following charitable foundations for their generous support to J.H: Wong Chun Hong Endowed Chair Professorship, Charlie Lee Charitable Foundation, and Fong Shu Fook Tong Foundation.

## Author contributions

J.H. and M.A. designed the experiments; K.K. and M.A. collected and analyzed the data for fiber photometry; K.K. and H.W. collected and analyzed the data for the behavioral part. K.K., G.Q., and L.Y. performed the surgeries and immunohistochemistry. M.A. wrote the manuscript. J.H. assisted in editing the manuscript. J.H. and M.A. supervised the project.

## Data Availability Statement

The data that support the findings of this study are available from the corresponding author upon reasonable request.

---

**Supplementary 1.**
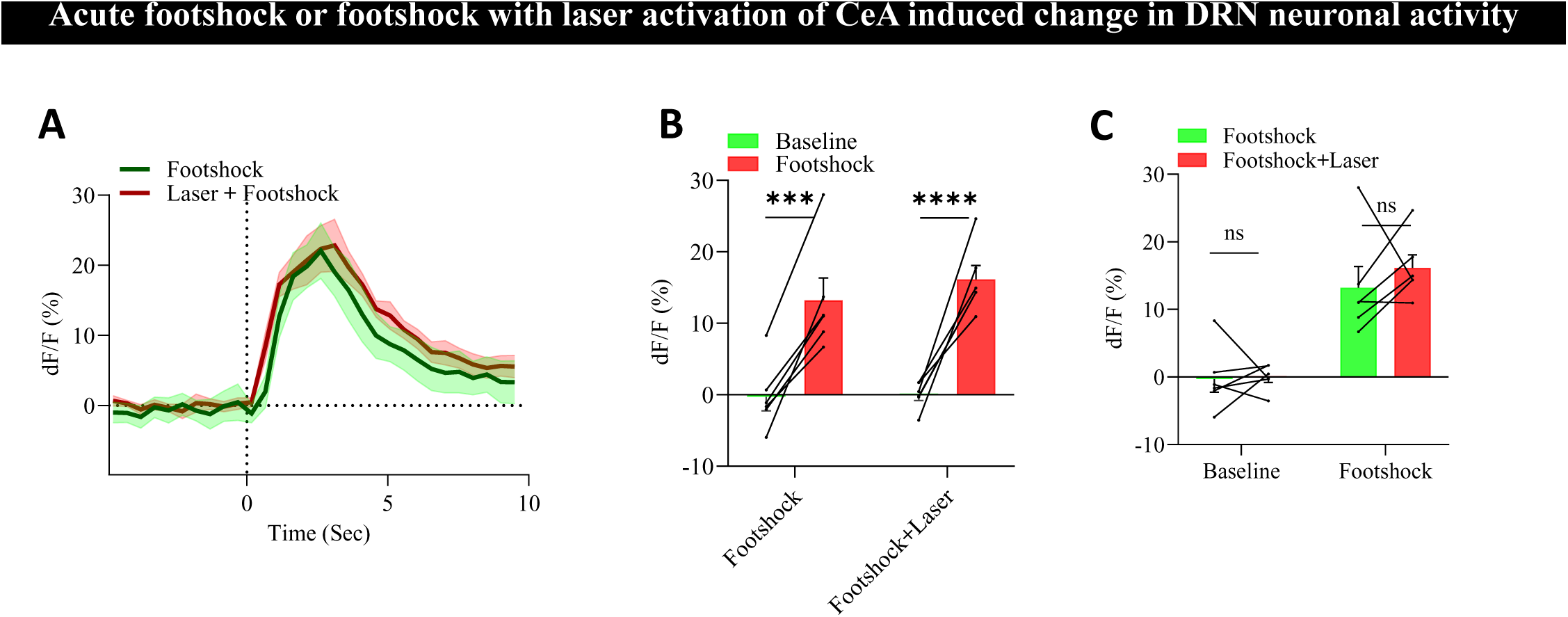
Acute foot shock and foot shock with laser activation of CeA induced change in DRN neuronal activity. **A.** Showing calcium traces during FS (aversive stimuli) and FS paired with CeA activation in mice. **B.** Average change in DRN activity before and during FS and FS paired with CeA activation in mice. RM Two-way ANOVA, Interaction: ^ns^P < 0.39, F (1, 10) = 0.79. Row Factor: ^ns^P < 0.55, F (1, 10) = 0.38. Column Factor: ****P < 0.0001, F (1, 10) = 97.17 Baseline vs FS: ***P < 0.0002, t = 6.34, df = 10. Baseline vs FS-Laser: ****P < 0.0001, t = 7.60, df = 10. Adjustment: Bonferroni. N = 6 mice. **C.** Showing the average changes between FS and FS paired with CeA activation in mice. RM Two-way ANOVA, Interaction: ^ns^P < 0.54, F (1, 10) = 0.40. Row Factor: ****P < 0.0001, F (1, 10) = 49.93. Column Factor: ^ns^P < 0.47, F (1, 10) = 0.57. Baseline (FS) vs Baseline (FS-Laser): ^ns^P > 0.9999, t = 0.085, df = 10. FS vs FS-Laser: ^ns^P > 0.9999, t = 0.98, df = 10. Adjustment: Bonferroni. N = 6 mice.

**Supplementary 2.**
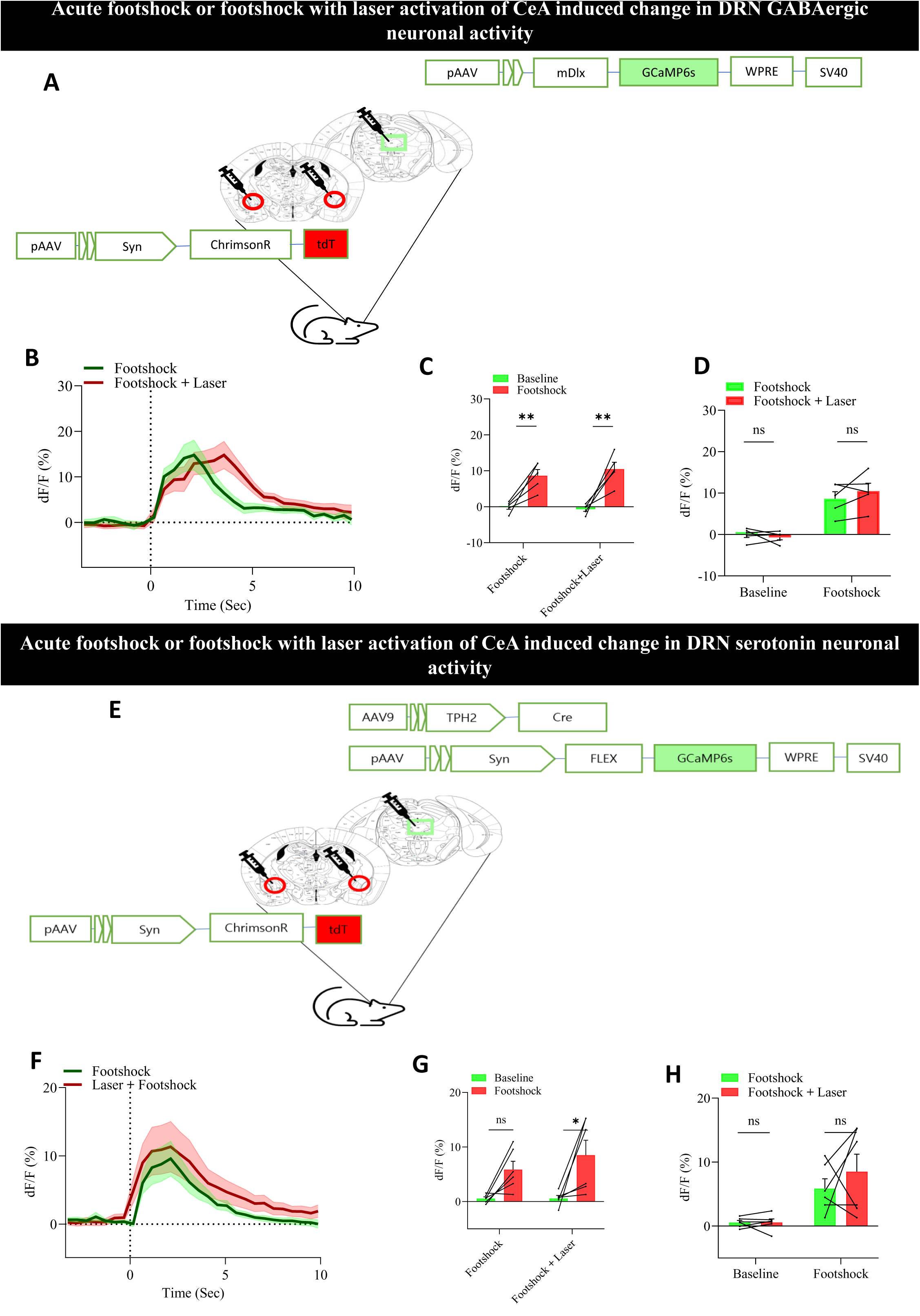
Acute foot shock or foot shock with laser activation of CeA induces a change in the subtype of DRN neuronal activity. **A.** Schematical representation of the experimental design for GCaMP6s virus injection. **B.** Showing calcium traces during FS (aversive stimuli) and FS paired with CeA activation in mDlx-GCaMP6s expression mice. **C.** Average change in DRN activity before and during FS and FS paired with CeA activation. RM Two-way ANOVA, Interaction: ^ns^P < 0.43, F (1, 8) = 0.67. Row Factor: ^ns^P < 0.62, F (1, 8) = 0.25 Column Factor: ***P < 0.0002, F (1, 8) = 42.71. Baseline vs FS: **P < 0.0075, t = 4.04, df = 8. Baseline vs FS-Laser: **P < 0.0016, t = 5.20, df = 8. Adjustment: Bonferroni. N = 5 mice. **D.** Showing the average changes between FS and FS paired with CeA activation. RM Two-way ANOVA, Interaction: ^ns^P < 0.17, F (1, 8) = 018. Row Factor: ***P < 0.0004, F (1, 8) = 32.79. Column Factor: ^ns^P < 0.49, F (1, 8) = 051. Baseline (FS) vs Baseline (FS-Laser): ^ns^P < 0.99, t = 0.53, df = 8. FS vs FS-Laser: ^ns^P < 0.32, t = 1.55, df = 8. Adjustment: Bonferroni. N = 5 mice. **E.** Schematical representation of the experimental design for GCaMP6s virus injection. **F.** Showing calcium traces during FS (aversive stimuli) and FS paired with CeA activation in TPH2-GCaMP6s expression mice. **G.** Average change in DRN activity before and during FS and FS paired with CeA activation. RM Two-way ANOVA, Interaction: ^ns^P < 0.44, F (1, 10) = 0.63. Row Factor: ^ns^P < 0.40, F (1, 10) = 0.75. Column Factor: **P < 0.0024, F (1, 10) = 16.30. Baseline vs FS: ^ns^P < 0.089, t = 2.29, df = 10. Baseline vs FS-Laser: *P < 0.013, t = 3.42, df = 10. Adjustment: Bonferroni. N = 6 mice. **H.** Showing the average changes between FS and FS paired with CeA activation. RM Two-way ANOVA, Interaction: ^ns^P < 0.44, F (1, 10) = 0.62. Row Factor: **P < 0.0015, F (1, 10) = 18.55. Column Factor: ^ns^P < 0.44, F (1, 10) = 0.65. Baseline (FS) vs Baseline (FS-Laser): ^ns^P < 0.99, t = 0.013, df = 10. FS vs FS-Laser: ^ns^P < 0.57, t = 1.133, df = 10. Adjustment: Bonferroni. N = 6 mice.

## References

Abdulla, Z. I., Mineur, Y. S., Crouse, R. B., Etherington, I. M., Yousuf, H., Na, J. J., & Picciotto, M. R. (2024). Medial prefrontal cortex acetylcholine signaling mediates the ability to learn an active avoidance response following learned helplessness training. Neuropsychopharmacology, 1–9.

Asim, M., Wang, H., Chen, X., & He, J. (2023). Potentiated GABAergic neuronal activities in the basolateral amygdala alleviate stress-induced depressive behaviors. CNS Neurosci Ther. 10.1111/cns.14422

Asim, M., Wang, H., Chen, X., & He, J. (2024). Potentiated GABAergic neuronal activities in the basolateral amygdala alleviate stressCinduced depressive behaviors. CNS Neuroscience & Therapeutics, 30(3), e14422.

Asim, M., Wang, H., & Waris, A. (2023). Altered neurotransmission in stress-induced depressive disorders: the underlying role of the amygdala in depression. Neuropeptides, 98, 102322.

Asim, M., Wang, H., Waris, A., & He, J. (2024). Basolateral amygdala parvalbumin and cholecystokinin-expressing GABAergic neurons modulate depressive and anxiety-like behaviors. Translational Psychiatry, 14(1), 418.

Babaev, O., Piletti Chatain, C., & Krueger-Burg, D. (2018). Inhibition in the amygdala anxiety circuitry. Experimental & molecular medicine, 50(4), 1–16.

Bartova, L., Dold, M., Kautzky, A., Fabbri, C., Spies, M., Serretti, A., Souery, D., Mendlewicz, J., Zohar, J., & Montgomery, S. (2019). Results of the European Group for the Study of Resistant Depression (GSRD)—basis for further research and clinical practice. The World Journal of Biological Psychiatry, 20(6), 427–448.

Bijata, M., Bączyńska, E., & Wlodarczyk, J. (2022). A chronic unpredictable stress protocol to model anhedonic and resilient behaviors in C57BL/6J mice. STAR protocols, 3(3), 101659.

Boukezzi, S., Costi, S., Shin, L. M., Kim-Schulze, S., Cathomas, F., Collins, A., Russo, S. J., Morris, L. S., & Murrough, J. W. (2022). Exaggerated amygdala response to threat and association with immune hyperactivity in depression. Brain, behavior, and immunity, 104, 205–212.

Cerniauskas, I., Winterer, J., de Jong, J. W., Lukacsovich, D., Yang, H., Khan, F., Peck, J. R., Obayashi, S. K., Lilascharoen, V., & Lim, B. K. (2019). Chronic stress induces activity, synaptic, and transcriptional remodeling of the lateral habenula associated with deficits in motivated behaviors. Neuron, 104(5), 899–915. e898.

Chourbaji, S., Zacher, C., Sanchis-Segura, C., Dormann, C., Vollmayr, B., & Gass, P. (2005). Learned helplessness: validity and reliability of depressive-like states in mice. Brain research protocols, 16(1-3), 70–78.

Cleare, A. J. (1997). Reduced whole blood serotonin in major depression. Depression and anxiety, 5(2), 108–111.

Clevenger, S. S., Malhotra, D., Dang, J., Vanle, B., & IsHak, W. W. (2018). The role of selective serotonin reuptake inhibitors in preventing relapse of major depressive disorder. Therapeutic advances in psychopharmacology, 8(1), 49–58.

Dale, E., Pehrson, A. L., Jeyarajah, T., Li, Y., Leiser, S. C., Smagin, G., Olsen, C. K., & Sanchez, C. (2016). Effects of serotonin in the hippocampus: how SSRIs and multimodal antidepressants might regulate pyramidal cell function. CNS spectrums, 21(2), 143–161.

Golden, S. A., Covington III, H. E., Berton, O., & Russo, S. J. (2011). A standardized protocol for repeated social defeat stress in mice. Nature protocols, 6(8), 1183–1191.

Groenewold, N. A., Opmeer, E. M., de Jonge, P., Aleman, A., & Costafreda, S. G. (2013). Emotional valence modulates brain functional abnormalities in depression: evidence from a meta-analysis of fMRI studies. Neuroscience & Biobehavioral Reviews, 37(2), 152–163.

Hartley, N. D., Gaulden, A. D., Báldi, R., Winters, N. D., Salimando, G. J., Rosas-Vidal, L. E., Jameson, A., Winder, D. G., & Patel, S. (2019). Dynamic remodeling of a basolateral-to-central amygdala glutamatergic circuit across fear states. Nature neuroscience, 22(12), 2000–2012.

Heifets, B. D., Salgado, J. S., Taylor, M. D., Hoerbelt, P., Cardozo Pinto, D. F., Steinberg, E. E., Walsh, J. J., Sze, J. Y., & Malenka, R. C. (2019). Distinct neural mechanisms for the prosocial and rewarding properties of MDMA. Science translational medicine, 11(522), eaaw6435.

Hutton, C., Déry, N., Rosa, E., Lemon, J., Rollo, C., Boreham, D., Fahnestock, M., Decatanzaro, D., Wojtowicz, J., & Becker, S. (2015). Synergistic effects of diet and exercise on hippocampal function in chronically stressed mice. Neuroscience, 308, 180–193.

Kostanjšak, L., & Zdunić, D. (2017). The role of thrombocyte serotonin system and some thrombocyte characteristics in treatment of depressive patients with cardiovascular diseases. Archives of Psychiatry Research, 53(1), 33.

Lesch, K.-P., & Waider, J. (2012). Serotonin in the modulation of neural plasticity and networks: implications for neurodevelopmental disorders. Neuron, 76(1), 175–191.

Liu, W.-Z., Zhang, W.-H., Zheng, Z.-H., Zou, J.-X., Liu, X.-X., Huang, S.-H., You, W.-J., He, Y., Zhang, J.-Y., & Wang, X.-D. (2020). Identification of a prefrontal cortex-to-amygdala pathway for chronic stress-induced anxiety. Nature communications, 11(1), 2221.

Marek, R., Strobel, C., Bredy, T. W., & Sah, P. (2013). The amygdala and medial prefrontal cortex: partners in the fear circuit. The Journal of physiology, 591(10), 2381–2391.

Moncrieff, J., Cooper, R. E., Stockmann, T., Amendola, S., Hengartner, M. P., & Horowitz, M. A. (2023). The serotonin theory of depression: a systematic umbrella review of the evidence. Molecular psychiatry, 28(8), 3243–3256.

Namburi, P., Beyeler, A., Yorozu, S., Calhoon, G. G., Halbert, S. A., Wichmann, R., Holden, S. S., Mertens, K. L., Anahtar, M., & Felix-Ortiz, A. C. (2015). A circuit mechanism for differentiating positive and negative associations. Nature, 520(7549), 675–678.

Nazzi, S., Maddaloni, G., Pratelli, M., & Pasqualetti, M. (2019). Fluoxetine induces morphological rearrangements of serotonergic fibers in the hippocampus. ACS Chemical Neuroscience, 10(7), 3218–3224.

Nikolova, Y. S., Misquitta, K. A., Rocco, B. R., Prevot, T. D., Knodt, A. R., Ellegood, J., Voineskos, A. N., Lerch, J. P., Hariri, A. R., & Sibille, E. (2018). Shifting priorities: highly conserved behavioral and brain network adaptations to chronic stress across species. Translational psychiatry, 8(1), 26.

Obermanns, J., Krawczyk, E., Juckel, G., & Emons, B. (2021). Analysis of cytokine levels, T regulatory cells and serotonin content in patients with depression. European Journal of Neuroscience, 53(10), 3476–3489.

Paquelet, G. E., Carrion, K., Lacefield, C. O., Zhou, P., Hen, R., & Miller, B. R. (2022). Single-cell activity and network properties of dorsal raphe nucleus serotonin neurons during emotionally salient behaviors. Neuron, 110(16), 2664–2679. e2668.

Pawluski, J. L., Paravatou, R., Even, A., Cobraiville, G., Fillet, M., Kokras, N., Dalla, C., & Charlier, T. D. (2020). Effect of sertraline on central serotonin and hippocampal plasticity in pregnant and non-pregnant rats. Neuropharmacology, 166, 107950.

Planchez, B., Surget, A., & Belzung, C. (2019). Animal models of major depression: drawbacks and challenges. Journal of Neural Transmission, 126, 1383–1408.

Prakash, N., Stark, C. J., Keisler, M. N., Luo, L., Der-Avakian, A., & Dulcis, D. (2020). Serotonergic plasticity in the dorsal raphe nucleus characterizes susceptibility and resilience to anhedonia. Journal of Neuroscience, 40(3), 569–584.

Pretorius, E. (2004). Corticosteroids, depression and the role of serotonin. Reviews in the Neurosciences, 15(2), 109–116.

Qiao, J., Tao, S., Wang, X., Shi, J., Chen, Y., Tian, S., Yao, Z., & Lu, Q. (2020). Brain functional abnormalities in the amygdala subregions is associated with anxious depression. Journal of Affective Disorders, 276, 653–659.

Ren, J., Isakova, A., Friedmann, D., Zeng, J., Grutzner, S. M., Pun, A., Zhao, G. Q., Kolluru, S. S., Wang, R., & Lin, R. (2019). Single-cell transcriptomes and whole-brain projections of serotonin neurons in the mouse dorsal and median raphe nuclei. Elife, 8, e49424.

Rosenkranz, J. A., Venheim, E. R., & Padival, M. (2010). Chronic stress causes amygdala hyperexcitability in rodents. Biological psychiatry, 67(12), 1128–1136.

Runia, N., Bergfeld, I. O., de Kwaasteniet, B. P., Luigjes, J., van Laarhoven, J., Notten, P., Beute, G., van den Munckhof, P., Schuurman, R., & Denys, D. (2023). Deep brain stimulation normalizes amygdala responsivity in treatment-resistant depression. Molecular psychiatry, 28(6), 2500–2507.

Sanacora, G., Treccani, G., & Popoli, M. (2012). Towards a glutamate hypothesis of depression: an emerging frontier of neuropsychopharmacology for mood disorders. Neuropharmacology, 62(1), 63–77.

Santomauro, D. F., Herrera, A. M. M., Shadid, J., Zheng, P., Ashbaugh, C., Pigott, D. M., Abbafati, C., Adolph, C., Amlag, J. O., & Aravkin, A. Y. (2021). Global prevalence and burden of depressive and anxiety disorders in 204 countries and territories in 2020 due to the COVID-19 pandemic. The Lancet, 398(10312), 1700–1712.

Son, H., Yang, J. H., Kim, H. J., & Lee, D. K. (2019). A chronic immobilization stress protocol for inducing depression-like behavior in mice. JoVE (Journal of Visualized Experiments)(147), e59546.

Sudimac, S., Sale, V., & Kühn, S. (2022). How nature nurtures: Amygdala activity decreases as the result of a one-hour walk in nature. Molecular psychiatry, 27(11), 4446–4452.

Talaee, N., Azadvar, S., Khodadadi, S., Abbasi, N., Asli-Pashaki, Z. N., Mirabzadeh, Y., Kholghi, G., Akhondzadeh, S., & Vaseghi, S. (2024). Comparing the effect of fluoxetine, escitalopram, and sertraline, on the level of BDNF and depression in preclinical and clinical studies: a systematic review. European Journal of Clinical Pharmacology, 80(7), 983–1016.

Walsh, J. J., Llorach, P., Cardozo Pinto, D. F., Wenderski, W., Christoffel, D. J., Salgado, J. S., Heifets, B. D., Crabtree, G. R., & Malenka, R. C. (2021). Systemic enhancement of serotonin signaling reverses social deficits in multiple mouse models for ASD. Neuropsychopharmacology, 46(11), 2000–2010.

Ye, J., & Veinante, P. (2019). Cell-type specific parallel circuits in the bed nucleus of the stria terminalis and the central nucleus of the amygdala of the mouse. Brain Structure and Function, 224, 1067–1095.

